## Supplementary information for "Flexible graphene-based neurotechnology for high-precision deep brain mapping and neuromodulation in Parkinsonian rats"

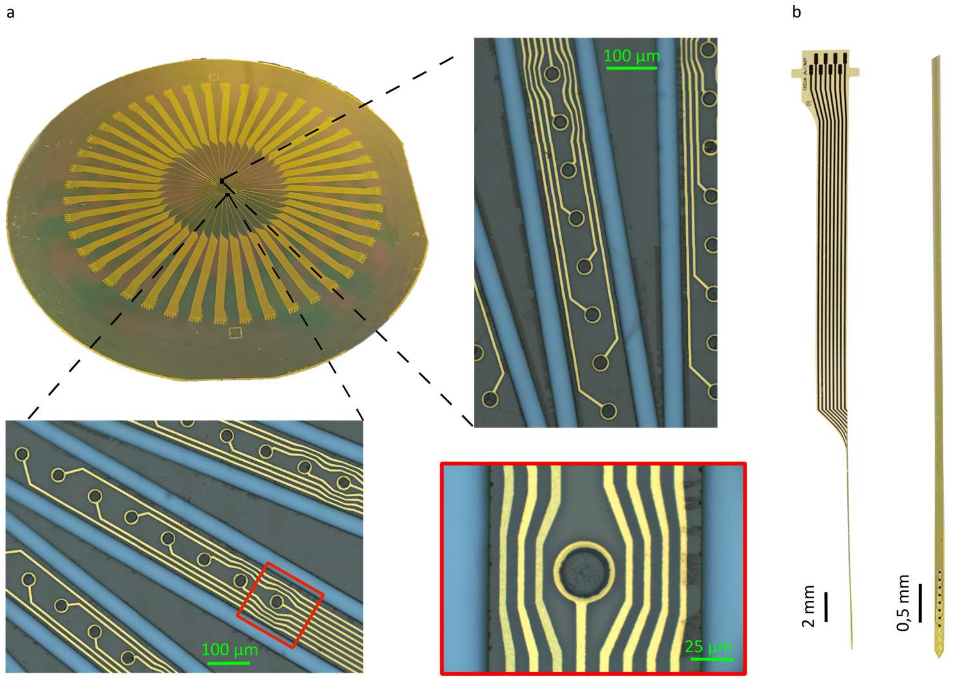


**Figure S1 | Flexible thin film microfabrication on a 4-inch Wafer. a**, Photograph of a 4-inch wafer with 44 devices microfabricated. 2 zooms of the central part allow for visualization of the tips of the leads where are placed the electrodes. With the red square, a zoom of a single rGO electrode. **b**, Photographs of a device peeled off from the wafer with a zoom of the thin tip.

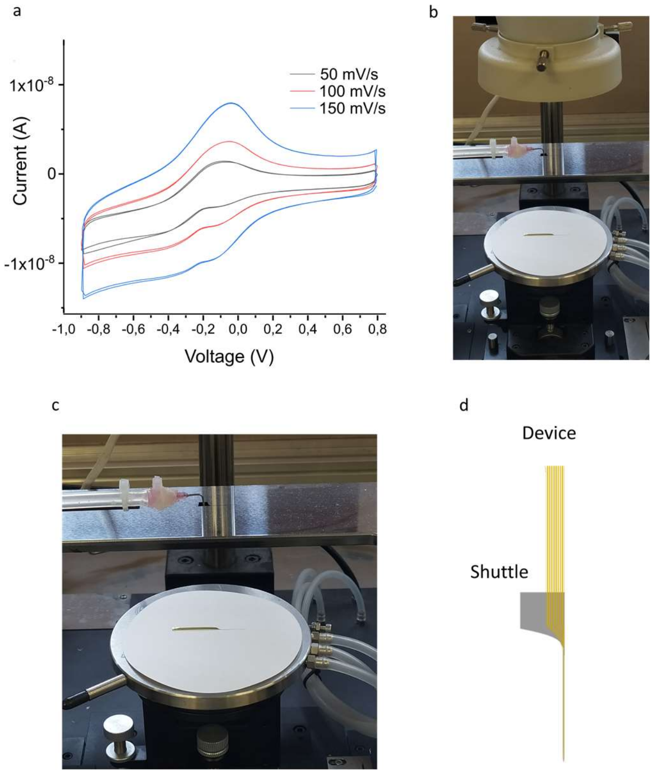


**Figure S2 | Electrochemical characterization and device attachment. a**, Cyclic voltammograms for one electrode were recorded at different scan rates, from -0.9 V to 0.8 V vs Ag/AgCl, in PBS. **b**,, Set-up used to glue with the adhesive polymer the flexible device to the shuttle. The device is placed on a probe station that can be moved with manipulators. A vacuum pump connects with a plastic tube a metallic tip, that can hold the shuttle. After applying the adhesive polymer with a brash on the tip of the shuttle, the flexible device is aligned with the shuttle looking under the microscope, and then assembled. **d,** Schematic of a device attached to the shuttle.

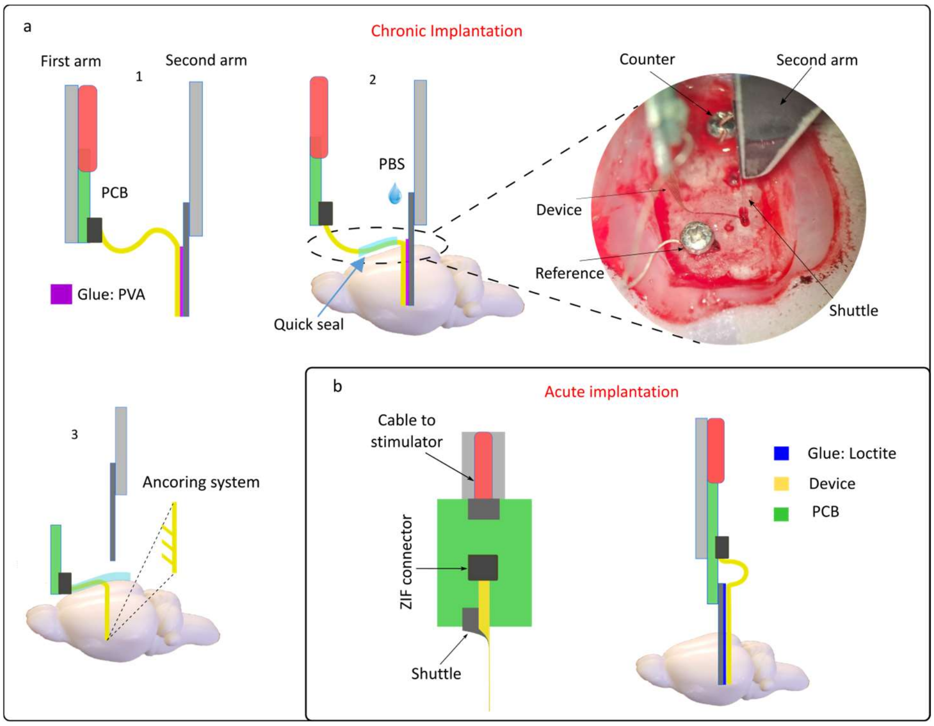


**Figure S3 | Insertion strategy in chronic and acute. a,** Chronic implantation with a 2 arms strategy in 3 steps. 1 The first arm holds the PCB that connects the tail of the lead, while the second one holds the shuttle attached to the array. 2 The second arm is slowly lowered with a micro driver to insert the device in the brain. When the insertion is completed, the tail is encapsulated with Kwik seal avoiding the insertion hall. PBS is then applied to dissolve the PVA that doesn't contact the brain fluids. The wires of the reference, ground and motor cortex connected to the PCB, are fixed on the scalp using small screws. 3 After 20 min, when the adhesive polymer is dissolved, the shuttle is removed and the device is kept in place thanks to the anchoring system. Everything is then encapsulated. **b**, Acute implantation with one arm insertion strategy; The shuttle is fixed in a free space of the PCB below the ZIF connector. The PCB is then connected to the stimulator and the cable is fixed on a stereotactic arm controlled by the micro driver. The device, glued with a permanent adhesive polymer, is slowly inserted in the brain until reaching the STN.

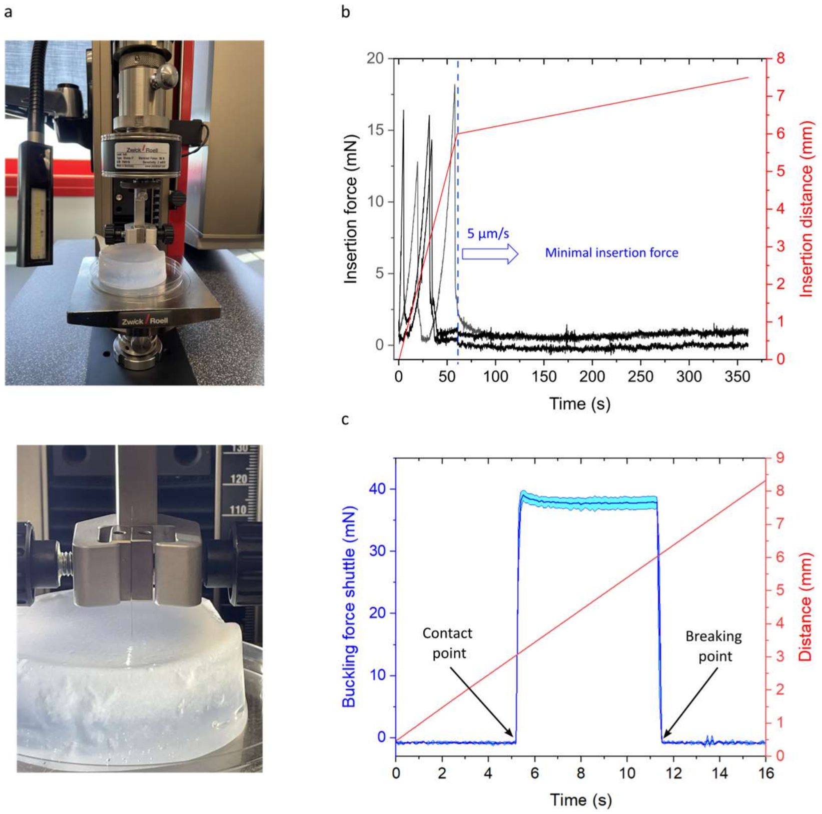


**Figure S4 | In vitro test of the insertion force. a,** Set-up for the measurement of the insertion force in the agar of a device glued with the shuttle. The device is inserted with the micro driver while a load force cell measures the force produced over time and the displacement . **b,** Insertion force (black) over time in agar for 3 devices glued with the shuttle, following the insertion speed used in vivo, 6 mm at 100 µm/s and

1.5 mm at 5 μm/s. Displacement over time in red. **c**, Average buckling force with STD for 3 shuttles on a rigid surface. The first point around 5 s represents the moment when the shuttle touches the rigid surface. Then it starts to bend under a force of around 37 mN, until the breaking point. The displacement is at the constant speed of 100 μm/s.

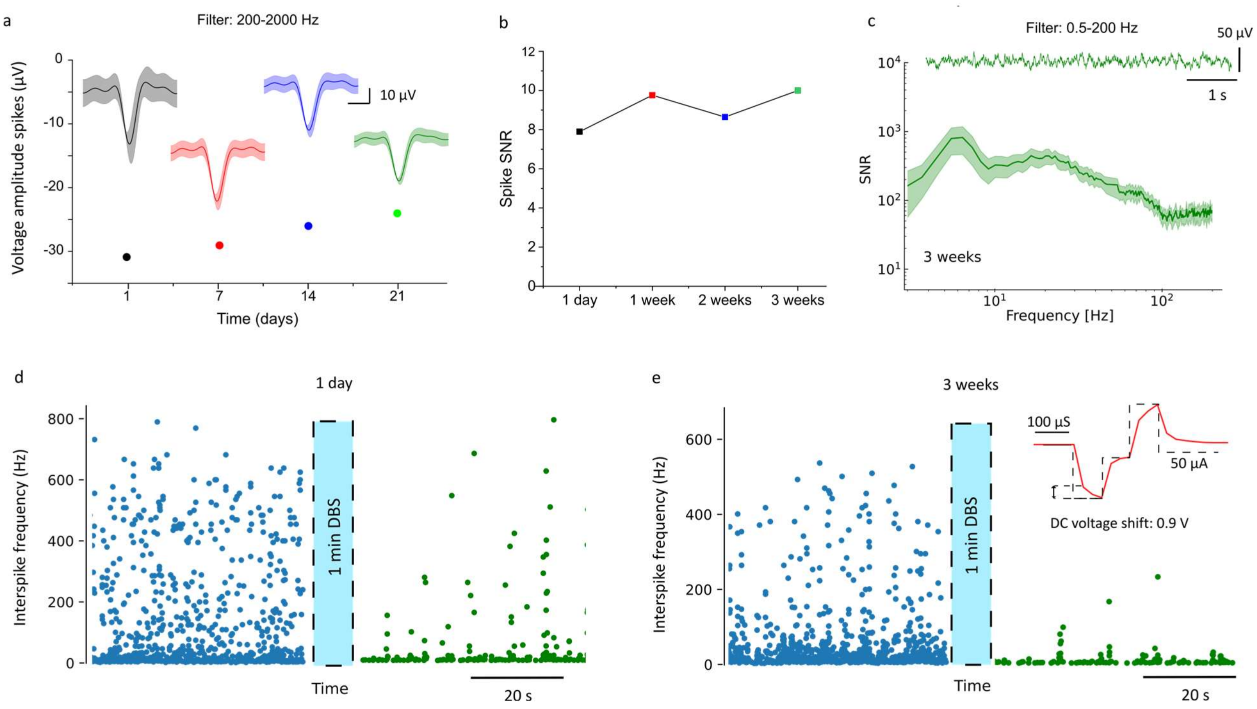


**Figure S5 | Proof-of-concept in vivo chronic study of recording and stimulation capabilities**. **a**, Average spike amplitude recorded over 3 weeks with one electrode in the STN. On top, the average voltage spike shapes with shadow area for the STD (over 100 s), are extracted at the corresponding time points. **b**, Spike to noise ratio over time of the experiment in the subpanel a. **c,** on top LFP recording at the third week with one electrode. Below is the average SNR spectrum with STD in the LFP range for the 8 electrodes of the array. **d**, **e**, IF distribution over time pre- (blue) and post-DBS (green), one day after implantation and after 3 weeks. In red, the measurement shows the electrode voltage polarization measured in the brain during the DBS protocol (biphasic pulses of 50 μA and 100 μs pulse width).

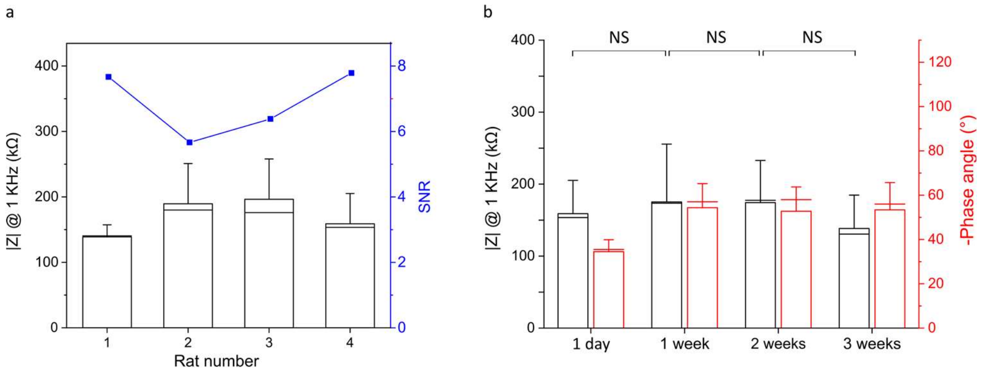


**Figure S6 | Electrodes in vivo chronic stability a**, **a**, Average impedance magnitude (black) at 1 kHz at day one of 4 arrays implanted in vivo, with the average spike-to-noise ratio (blue) of one recording electrode in the STN of each implant. **b**, Average impedance magnitude (black) and phase (red) of the 8 electrodes of one array measured in a chronically implanted rat over weeks (magnitude after 1 week NS p = 0.62, after 2 weeks NS p = 0.98, after 3 weeks NS p = 0.19, in the Welch T-test).

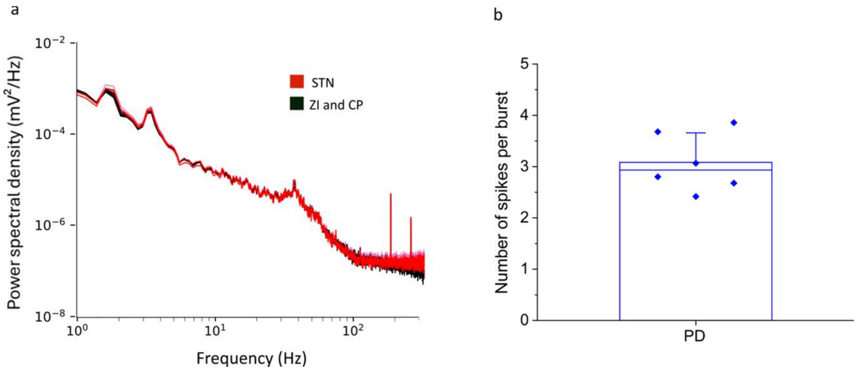


**Figure S7 | Power spectral density and burst length. a**, PSD through the electrodes of one array implanted in the STN. The red one represents the 2 electrodes in the STN while the black ones are placed in the Zona Incerta (ZI) and internal capsule (CP). **b**, Spike to noise ratio over time in the STN of one rat chronically implanted. **b**, Average number of spikes per burst recorded over 100 s in the STN of the PD group.

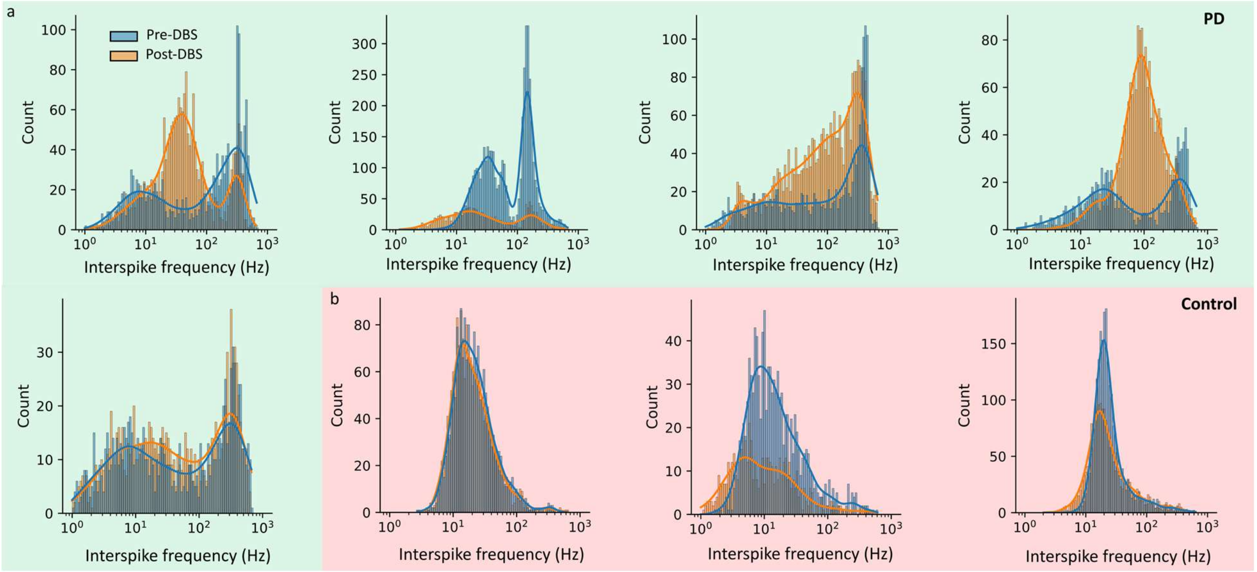


**Figure S8 | Spike distribution in the PD-Control. a,b** IF distribution pre- (blue) and post-DBS (orange) for the spikes recorded in the STN of 5 PD (green rectangle) and 3 control rats (red rectangle). The histograms are fitted with the kernel density probability **c**, Fano factor quantified over 100 s pre-DBS in the MUA range (200-2000 Hz) in the STN of the 2 groups, PD and Control (*p =0,039 in the Welch T-test) **d**, Average number of spikes per burst recorded over 100 s in the STN of the PD group.

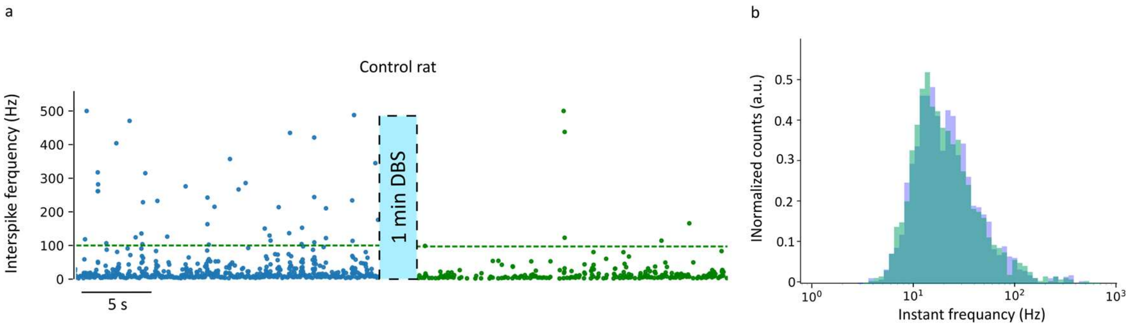


**Figure S9 | Effects of DBS on control rat. a**, Interspike frequency in the STN of a control rat, pre (blue) and post (green) DBS (75 µA at 130Hz). The green dash line highlights the cut-off frequency of the IF for the burst and tonic events. **b**, Normalized interspike frequency distribution over 100 s for the same control rat, pre and post-DBS.µ

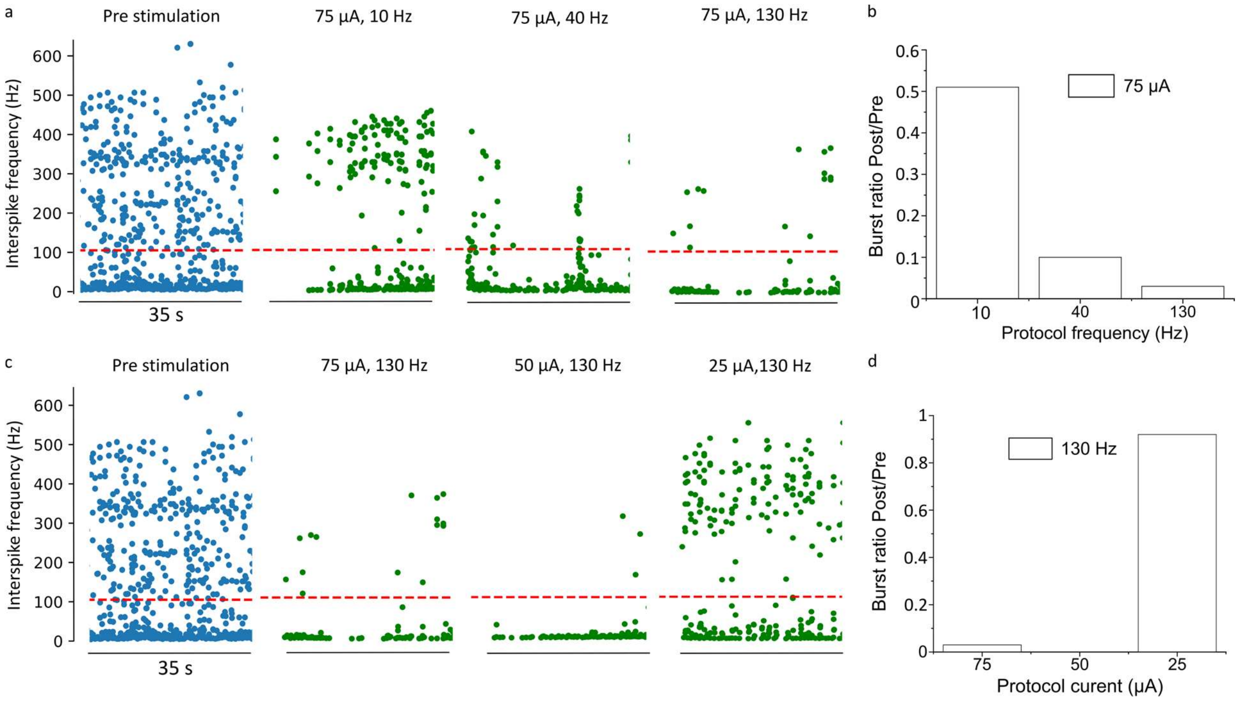


**Figure S10 | Effects of DBS parameters on STN spiking activity**. **a**, IF over time in the STN of a PD rat in period pre- (blue) and post-DBS (green), for biphasic current pulses of 100 µs and an amplitude of 75uA. Different stimulation frequencies were tested, 10Hz, 40Hz and 130 Hz. The red dash line highlights the cut-off frequency usually used to distinguish burst and tonic IF. **b,** Quantification over 35 s of the burst ratio post/pre for the 3 DBS protocols. **c**, The same effects are evaluated at a fixed frequency, 130 Hz and variable current, 25, 50 and 75 µA. **d**, Quantification over 35 s of the burst ratio post/pre for the 3 DBS protocols of variable current.

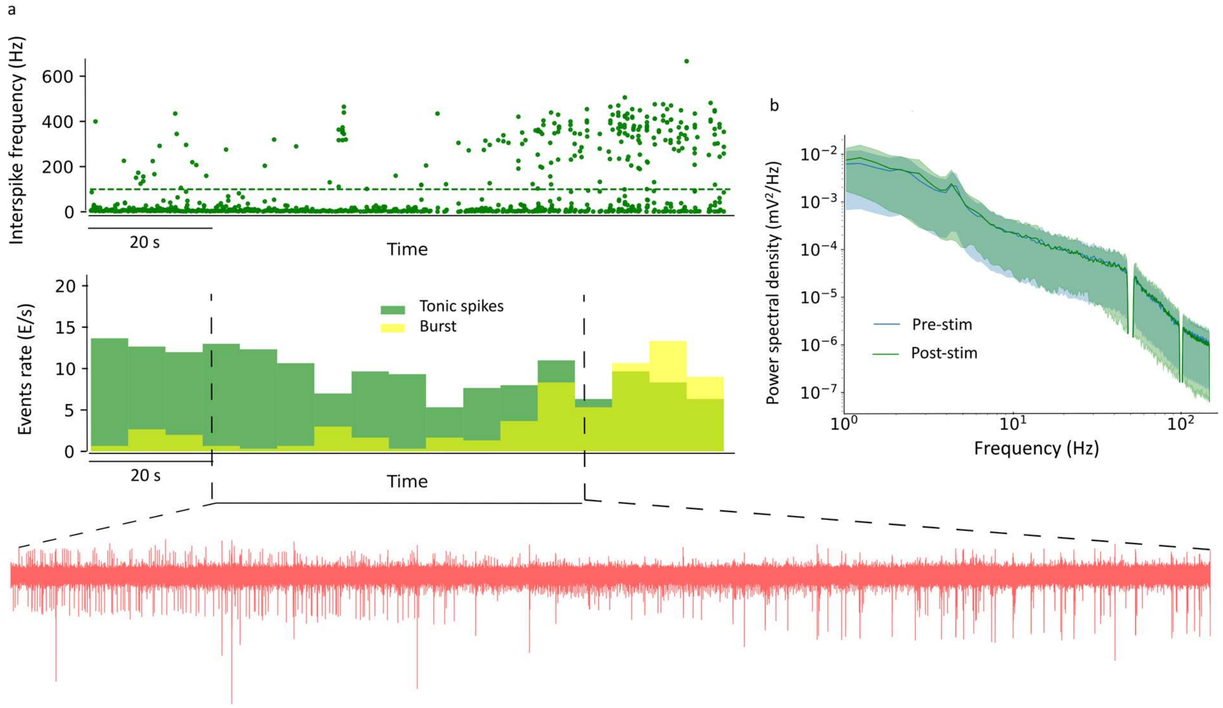


**Figure S11 | STN Spiking activity evolution post-DBS**. **a,** IF over time in the STN of a PD rat several minutes after DBS (75 µA, 130 Hz). The histogram below quantifies the burst rate (yellow) and tonic spike rate (green) in the same period in 6 s bins. Below is the corresponding zoom of the recording between the 2 dashed lines in the spike range (200-2000 Hz) to visualize the switch from tonic to burst activity. **b**, Average PSD in the STN over 100 s with shadow area for the STD of the PD group, pre-DBS in blue and post-DBS in green.
